# An Effective mRNA-LNP Vaccine Against the Lethal Plague Bacterium

**DOI:** 10.1101/2022.08.07.503096

**Authors:** Edo Kon, Yinon Levy, Uri Elia, Hila Cohen, Inbal Hazan-Halevy, Moshe Aftalion, Assaf Ezra, Erez Bar-Haim, Gonna Somu Naidu, Yael Diesendruck, Shahar Rotem, Nitay Ad-El, Meir Goldsmith, Emanuelle Mamroud, Dan Peer, Ofer Cohen

**Affiliations:** Laboratory of Precision NanoMedicine, Shmunis School for Biomedicine and Cancer Research, George S. Wise Faculty of Life Sciences, Tel Aviv University, Tel Aviv 69978, Israel; Center for Nanoscience and Nanotechnology, Tel Aviv University, Tel Aviv 69978, Israel; Department of Materials Sciences and Engineering, Iby and Aladar Fleischman Faculty of Engineering, Tel Aviv University, Tel Aviv 69978, Israel; Cancer Biology Research Center, Tel Aviv University, Tel Aviv 69978, Israel; Department of Biochemistry and Molecular Genetics, Israel Institute for Biological Research, Ness-Ziona 76100, Israel

**Author notes:** These authors contributed equally to this work (Edo Kon, Yinon Levy, Uri Elia). These authors contributed equally to this work (Emanuelle Mamroud, Dan Peer, Ofer Cohen).

## Abstract

Plague is a rapidly deteriorating contagious disease that has killed millions of people during the history of mankind and is caused by the gram-negative bacterium *Yersinia pestis*. Currently, the disease is treated effectively with antibiotics. However, in the case of an outbreak caused by a multiple-antibiotic-resistant strain, alternative countermeasures are required. Despite the many efforts to develop a safe vaccine against the disease, there is still no vaccine approved for use in western countries. mRNA Lipid Nanoparticle (mRNA-LNP) vaccines have been demonstrated during the Covid-19 pandemic to be a versatile, clinically relevant, and rapidly manufactured vaccine platform. However, harnessing this platform for bacterial pathogens remains a formidable challenge. Here, we describe the design of several mRNA-LNP vaccine versions against *Y. pestis*, based on the F1 capsular antigen. We demonstrate that mRNA-LNP vaccines encoding the F1 antigen with either no signal sequences or conjugated to human Fc, provide substantial cellular and humoral responses. Most importantly, these vaccine candidates fully protect animals against *Y. pestis* infection. The results of this study suggest that mRNA-LNPs can be effective as anti-bacterial vaccines, and further developed to combat other bacterial pathogens, which are urgently needed, given the looming threat of antibiotic resistance.

**One-Sentence Summary:** A novel mRNA-LNP vaccine against *Y. pestis*, the etiological agent of plague and the first documented mRNA-LNP vaccine to protect against a lethal bacterial pathogen infection.

## Main Text

Plague is an infectious disease caused by the gram-negative bacterium *Yersinia pestis (Y. pestis)*, which has claimed the lives of millions of people throughout human history in three major pandemics and numerous local outbreaks (*1*). Due to its lethality and infectivity, *Y. pestis* is classified as a potential bio-terror agent (*2*). Morbidity and mortality from plague have decreased significantly since the introduction of antimicrobials, but the isolation of *Y. pestis* strains resistant to multiple therapeutic antibiotic (*3, 4*), and the fear of a natural or intentional disease outbreak caused by antibiotic-resistant strains emphasize the need to develop vaccines against this deadly disease. Much effort has been invested in recent decades by research groups around the world to develop a safe and efficacious vaccine against plague (*5*). Although several candidate vaccines were found to confer protection in animal models of plague and in clinical studies, none of the candidates was approved for use against plague in western countries (*6*). Sub-unit vaccines against plague are based on the two major protective antigens; the low-calcium response virulence (LcrV) protein and the F1 capsule antigen (*5*). We have recently demonstrated that recombinant F1 induces rapid onset of protective immunity in plague mouse models and thus could serve as a vaccine for emergency situations such as an outbreak involving antibiotic-resistant strains (*7*).

Recently, mRNA lipid nanoparticles (LNPs) vaccines have been recognized as a breakthrough vaccine platform as a result of their effectiveness in the fight against the COVID-19 pandemic (*8*). However, while a multitude of mRNA vaccines are designed as viral or cancer vaccines, bacterial pathogens remain largely untapped and are extremely important due to the emerging antibiotic-resistance crisis (*9, 10*). mRNA-LNPs can serve as an important tool in the arsenal towards this goal since they are clinically relevant, rapidly manufactured and inherently modular. Very few studies to date have addressed the implementation of the mRNA vaccine platform against bacterial pathogens, mostly reporting modest protection or decrease in bacterial burden (*11-14*).

Our study aimed to harness the clinically applicable mRNA-LNP vaccine platform as a means of designing an effective vaccine against this lethal bacterial pathogen. Based on the F1 antigen, we designed several vaccine versions, evaluated the antigen-specific humoral and cellular responses, and determined protection effectiveness in mice challenged with a fully virulent *Y. pestis* strain.

First, we designed an mRNA construct coding for the bacterial F1 capsular antigen (Caf1). Vaccination with the F1 protein was shown to effectively induce anti-F1 antibodies and provide protection against bubonic plague in murine models (*7, 15, 16*). In the current study, we aimed to evaluate the immunogenicity of an mRNA vaccine platform encoding the F1 protein, and thus rationalized that to achieve activation of a humoral immune response, the encoded protein should be directed towards the secretory pathway. Therefore, a signal peptide (SP) sequence originating from the human Ig light chain was introduced upstream to the *caf1* gene, replacing the native bacterial signal sequence, resulting in the SP-*caf1* mRNA construct (Fig. 1A). The mRNA was encapsulated in a previously reported LNP formulation that we demonstrated to be highly effective as an mRNA-LNP vaccine (*17, 18*) (Fig. 1B,C). Physicochemical characterization of SP-*caf1* mRNA-LNPs resulted in an average size of 61.5 nm, and a PDI of 0.132 (Fig. 1D), and the uniform size distribution was corroborated by cryogenic electron microscopy (Cryo-EM) analysis (Fig. 1E).

**Fig. 1.**
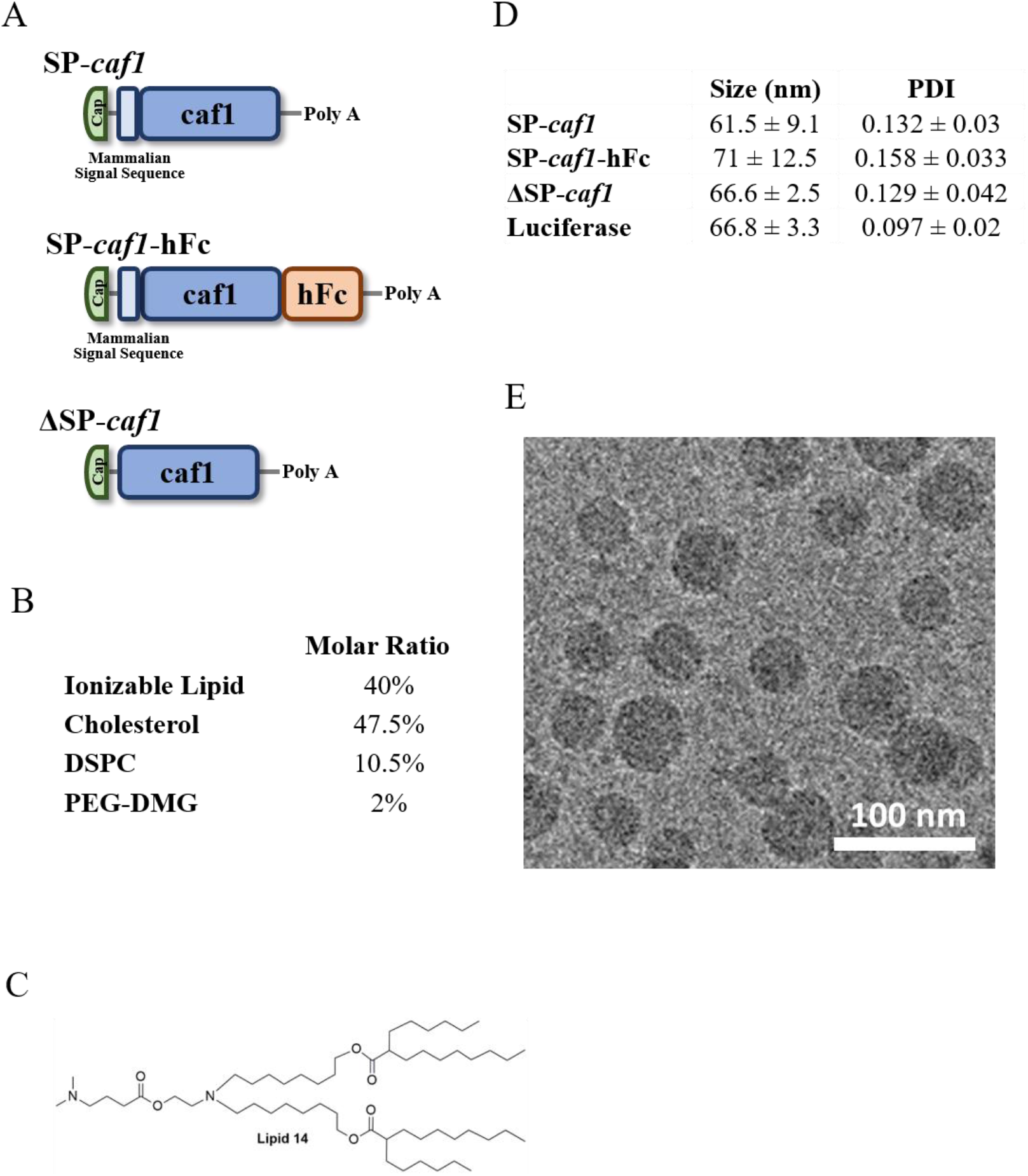
Construct design and physicochemical characterization of mRNA-LNPs formulations used throughout the study. (A) Schematic representation of the mRNA constructs used throughout the study: SP-*caf1*, SP-*caf1*-hFc and ΔSP-*caf1*. (B) LNP formulation utilized throughout the study. (C) Chemical structure of the proprietary ionizable lipid – Lipid 14. (D) Table summarizing LNP physicochemical aspects. (E) A representative Cryo-EM image of LNP-encapsulated SP-*caf1* mRNA. Scale bar 100 nm

We determined the expression of the F1 protein in both supernatant and cell pellets and confirmed that the protein is expressed and secreted (Fig. 2A). Next, we assessed the *in-vivo* immunogenicity of the SP-*caf1* mRNA-LNPs. Mice were intramuscularly immunized thrice, three weeks apart, with 5 µg of SP-*caf1* mRNA-LNPs (Fig. 2B). A robust and statistically significant cellular immune response was determined in both SP-*caf1* mRNA-LNPs and recombinant F1 (rF1)-vaccinated mice, two weeks after administration of the third vaccine dose. In contrast, humoral response was recorded only in rF1-vaccinated mice (Fig. 2C,D). Supported by previous studies conducted in our lab which demonstrated that animals that do not develop anti-F1 antibody titers succumb to bubonic plague infection (*7*), we devised two approaches to elicit a protective humoral response. First, we designed a human Fc-conjugated SP-*caf1* (SP-*caf1*-hFc) (Fig. 1A), based on the rational that hFc conjugation was reported to increase stability, half-life and immunogenicity of the targeted protein (*19-21*). Next, we designed an mRNA construct encoding a non-secreted F1 antigen by removing the mammalian signal peptide from the mRNA sequence (ΔSP-*caf1*) (Fig. 1A). Since signal peptides direct the expressed proteins to the secretory pathway, i.e. to the endoplasmic reticulum (ER) and Golgi compartments, where a variety of post-translational modifications (PTMs) naturally occur (*21*), we hypothesized that the intracellular mammalian translation of bacterial antigens which include a signal peptide may yield proteins which are different, and might affect the immunogenicity, compared to proteins naturally produced by the bacteria itself. The physicochemical properties of the two mRNA-LNPs formulations can be seen in Fig. 1D.

**Fig. 2.**
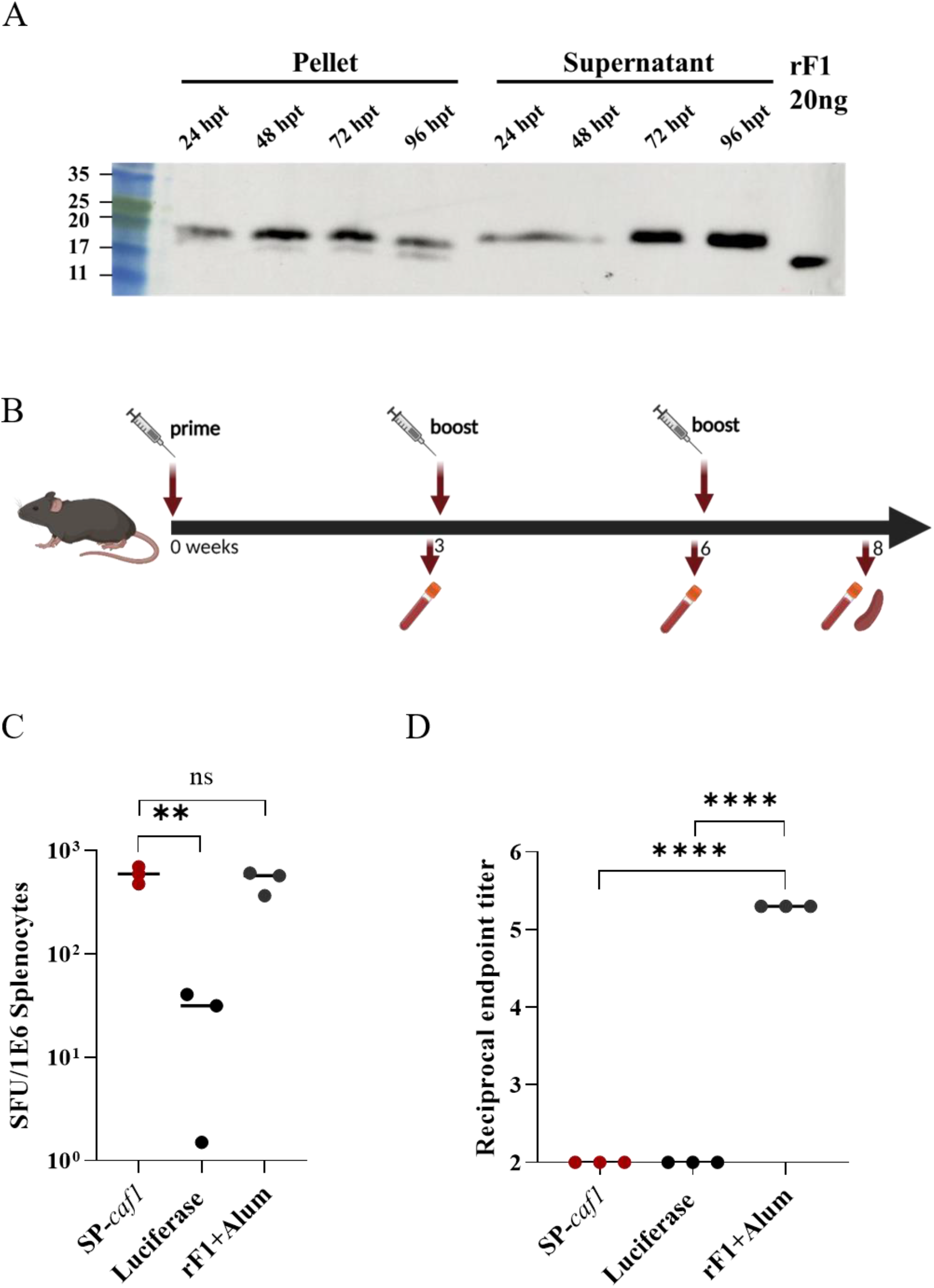
SP-*caf1* mRNA-LNPs induce potent cellular but not humoral immune responses. (**A**) Western blot analysis of F1 expression in HeLa cells transfected with SP-*caf1* mRNA-LNPs (2 μg/mL, 24−96 h). Purified recombinant F1 (20ng) was used as reference for protein migration (**B**) Schematic diagram of immunization and sample collection. C57BL/6 mice were immunized with 5µg SP-*caf1* or Luciferase mRNA-LNPs or with 80µg recombinant F1 protein with alum and boosted with an equivalent dose 21 and 42 days later. Serum samples were collected at days 20, 41 and 56. Spleens were collected at day 63. Illustration created with BioRender.com. (**C**) F1-specific cellular response determined by ELISpot. Splenocytes from vaccinated animals were stimulated with 10µg/ml F1 peptide mix for 24hrs, and the frequency of IFNγ-secreting cells was determined using the ELISpot assay. (**D**) Anti-F1 IgG titers were detected from serum samples at the indicated time points via an ELISA assay (n=3). Statistical analysis was performed using a one-way ANOVA followed by post hoc Newman–Keuls test (**, *p* < 0.01; ***, *p* < 0.001; ****, *p* < 0.0001).

Thereafter, we aimed to evaluate the immune responses elicited by SP-*caf1*-hFc and ΔSP-*caf1*-based mRNA-LNP vaccines. SP-*caf1*-hFc and ΔSP-*caf1* mRNAs were encapsulated in the lipid formulation previously described (Fig.1 B,C) and protein expression was confirmed (Fig. 3A). As expected, F1 expressed from the SP-*caf1*-hFc mRNA construct was detected mostly in the supernatant fraction while F1 encoded by ΔSP-*caf1* was expressed predominantly in the cellular lysate fraction, confirming the inability of the protein to enter the secretory pathway (Fig. 3A).

**Fig. 3.**
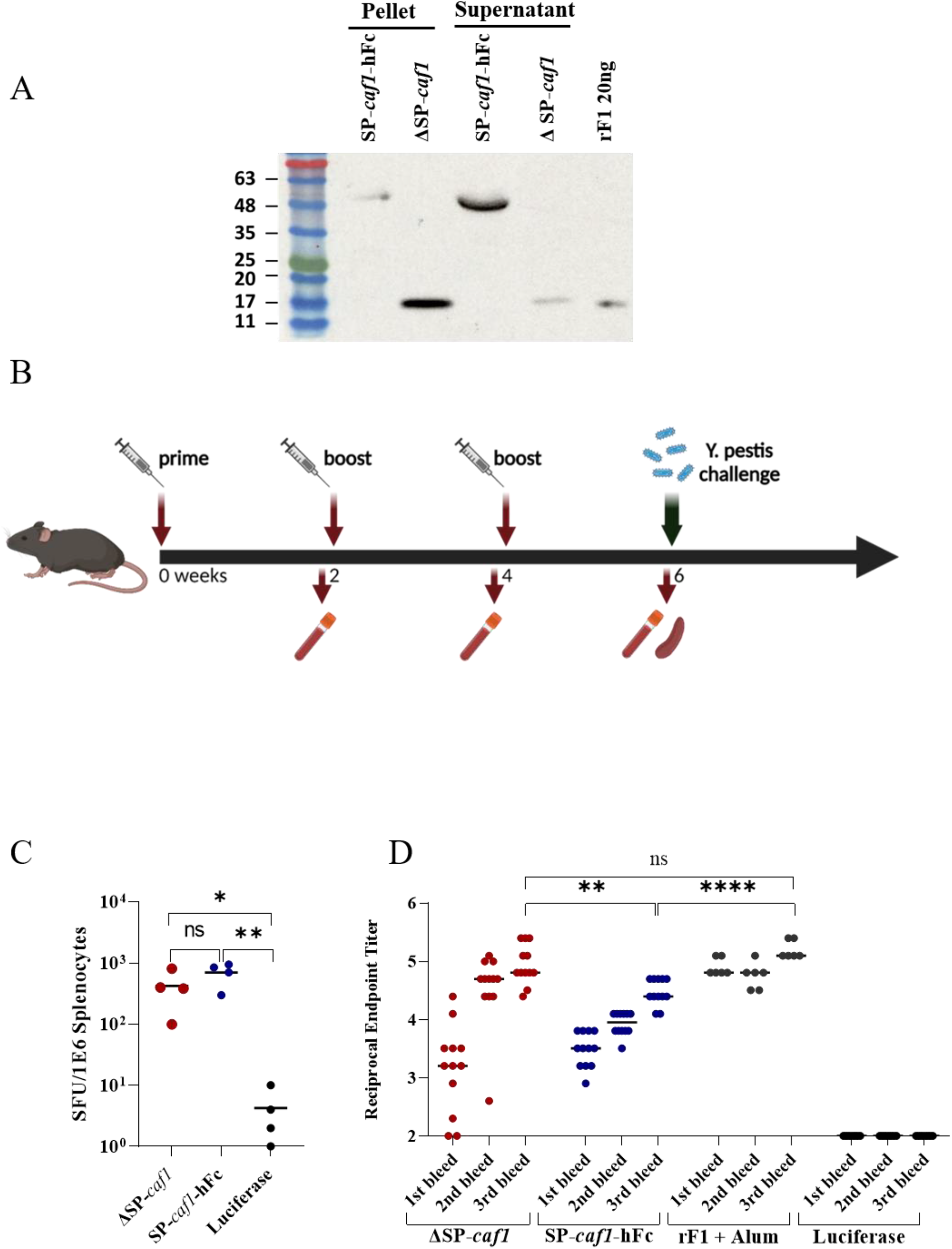
SP-*caf1*-hFc and ΔSP-*caf1* mRNA LNPs elicit robust cellular and humoral immune responses. (**A**) Western blot analysis of HeLa cells transfected with SP-*caf1*-hFc and ΔSP-*caf1* mRNA-LNPs (0.5 μg/mL, 72 h). (**B**) Schematic diagram of immunization and sample collection. C57BL/6 mice were immunized with mRNA-LNPs encapsulating SP-*caf1*-hFc, ΔSP-*caf1* or luciferase (5µg) or with 80µg recombinant F1 protein with alum and boosted with an equivalent dose 14 and 28 days later. Serum samples were collected at days 13, 27 and 42. Spleens were collected at day 42. Illustration created with BioRender.com. (**C**) F1-specific cellular response determined by ELISpot. Splenocytes from vaccinated animals were stimulated with 10µg/ml F1 peptide mix for 24hrs, and the frequency of IFNγ-secreting cells was determined using the ELISpot assay (n=4). (**D**) Anti-F1 IgG titers were detected from serum samples at the indicated time points via an ELISA assay (n=12). Statistical analysis was performed using a one-way ANOVA followed by post hoc Newman–Keuls test (*, *p* < 0.05; **, *p* < 0.01; ***, *p* < 0.001; ****, *p* < 0.0001).

To evaluate the *in-vivo* immunogenicity of F1 expressed from SP-*caf1*-hFc and ΔSP-*caf1* mRNA-LNPs, mice were immunized thrice, two weeks apart, with 5 µg of each construct (Fig 3B). Antigen-specific cellular responses were recorded following vaccination with both SP-*caf1*-hFc and ΔSP-*caf1* mRNA-LNPs (Fig. 3C), similar to the original SP-*caf1* construct. However, immunization with both SP-*caf1*-hFc and ΔSP-*caf1* mRNA-LNPs led to a robust, time-dependent anti-F1 humoral response which was comparable to that elicited by rF1 adjuvanted with alum (rF1+Alum) (Fig. 3D).

Given the high levels of F1-binding antibodies and strong cellular response recorded in both ΔSP-*caf1* and SP-*caf1*-hFc mRNA-LNP immunized-mice, we hypothesized that the vaccinated animals would be able to withstand a lethal *Y. pestis* challenge in the bubonic plague mouse model. Consequently, all vaccinated animals were inoculated subcutaneously with a lethal dose of the fully virulent *Y. pestis* strain Kimberley53 (Kim53). Encouragingly, both ΔSP-*caf1* and SP-*caf1*-hFc mRNA-LNPs immunizations were able to confer full protection against the lethal challenge (Fig. 4). In accordance with previous reports (*15, 23*), all control mice succumbed to the infection by day 7 whereas all rF1+alum-vaccinated animals survived the lethal challenge.

**Fig. 4.**
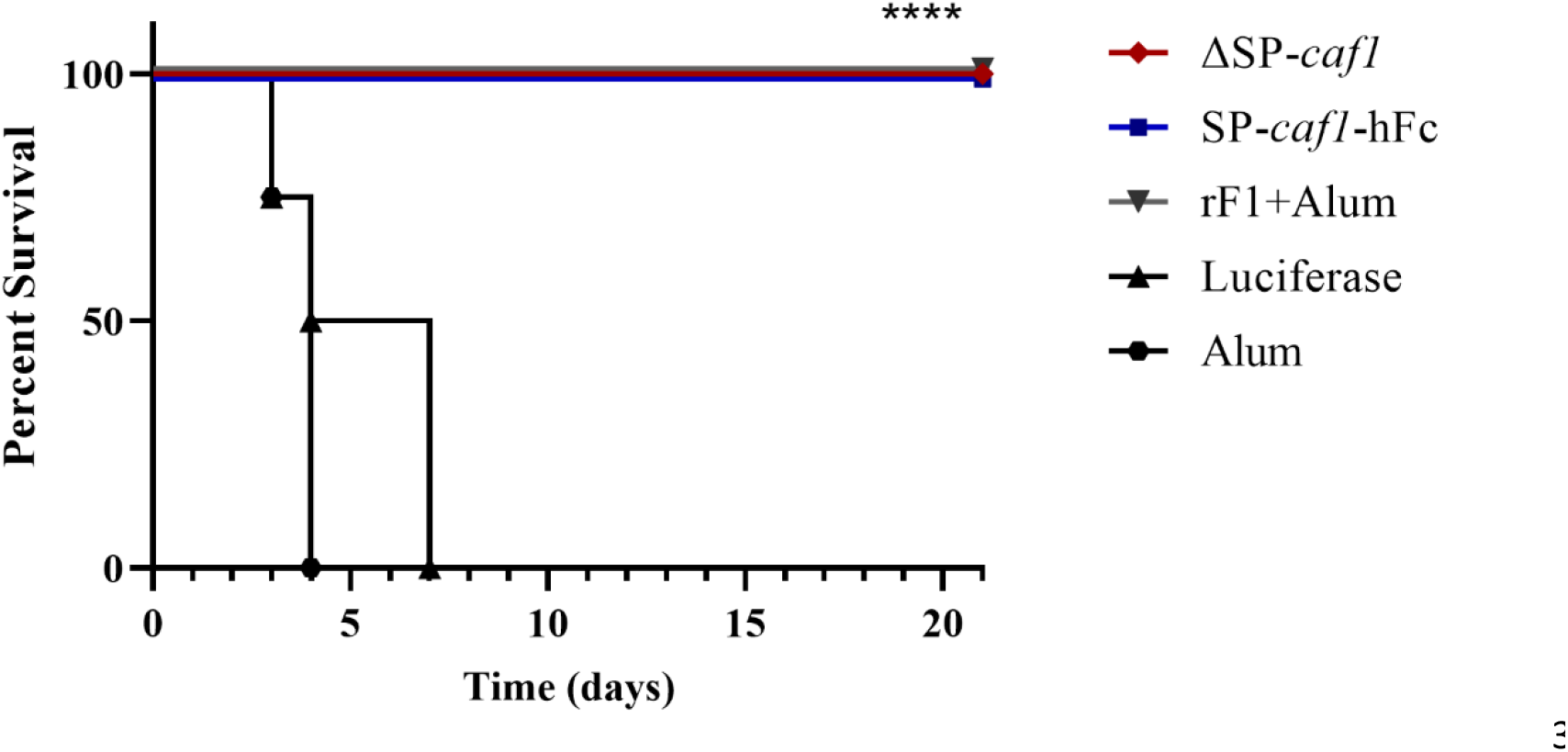
ΔSP-*caf1* and SP-*caf1*-hFc mRNA-LNPs protect mice against a lethal *Y. pestis* challenge in the bubonic plague model. C57BL/6 mice were immunized with SP-*caf1*-hFc, ΔSP-*caf1* mRNA-LNPs (5µg) (n=8), luciferase mRNA-LNPs (5µg) (n=4), recombinant F1 protein (80µg) with alum (n=6) or alum alone (n=4) and boosted with an equivalent dose 14 and 28 days later. 14 days following the last boost, mice were inoculated subcutaneously with the fully virulent *Y. pestis* strain Kim53 (100cfu), and monitored for two weeks following the challenge. Statistical analysis was performed using the log-rank (Mantel−Cox) test (****, *p* < 0.0001).

Here we describe the successful utilization of an mRNA-LNP vaccine platform for protection against the notorious bacterial pathogen *Y. pestis*, that causes plague. In contrast to viral proteins, which are naturally adapted for expression by eukaryotic systems, the construction of an mRNA vaccine platform that encodes for the expression of a bacterial protein requires adaptations. It is therefore challenging to design mRNA vaccines that are effective against bacterial pathogens. Thus, to enable the efficient expression, translocation and secretion of the F1 outer-membrane protein by eukaryotic systems, we replaced the F1 prokaryotic secretion signal with a eukaryotic one (Fig. 1A). In the natural *Y. pestis* expression environment, the newly synthesized F1 protein is translocated to the periplasm of the bacterial cell, and transported to the cell surface with the assistance of a chaperon and an anchoring protein, resulting in high-molecular-weight F1 oligomers (*24*). Since these elements do not exist in the eukaryotic expression system, we expressed F1 as a monomer by using the circular permutated form of the protein (*16*).

The secreted *caf1* mRNA-LNP vaccine yielded antigen-specific cellular responses but not humoral responses. While cellular responses can be beneficial for protection against bacterial pathogens with a predominantly intra-cellular life-cycle, such as *Mycobacterium tuberculosis* (*25*), a parallel robust humoral response is required for bacteria in which life cycle is mainly extra-cellular.

The current study demonstrates that both Fc-conjugated and signal peptide-devoid *caf1* mRNA sequences induce powerful antigen-specific humoral and cellular responses that protect mice from a lethal *Y. pestis* challenge. However, the SP-caf1 mRNA sequence induced only cellular responses. It is possible that this phenomenon is due to differences in intracellular trafficking between secreted and non-secreted bacterial proteins. For example, post-translational modifications such as glycosylation can occur in the secretory pathway and were reported to impair immune responses to nucleic acid based vaccines (*26*). Alternatively, and as we here demonstrate, factors such as serum stability, which is conferred by the Fc conjugation may contribute to the improved immune response, even when the expressed protein is shuttled via secretory pathways. Therefore, the characterization of the exact mechanisms that alter immunogenicity to F1 in the secretory pathway and the mechanism leading to the strong humoral responses to a non-secreted antigen are of great interest and remain to be elucidated.

The data presented in the current study suggests that the mRNA-LNP platform can be harnessed for effective vaccination against bacterial pathogens. These findings are of great relevance and immense importance, in light of the global emerging crisis of antibiotic resistance and the lack of efficient conventional therapies and vaccine candidates.

## Acknowledgments

EK thanks the Yoran Institute for Human Genome Research for their support. We thank Dr. Ronit Rosenfeld for helpful discussions and Dr. Ohad Mazor and Dr. Shmuel Yitzhaki for support throughout the study We thank the Biotechnology Department at IIBR for providing us the recombinant F1 protein. We thank Dr. Inbal Abutbul Ionita and Prof. Dganit Danino from the CryoEM Laboratory of Soft Matter, Faculty of Biotechnology and Food Engineering, Technion-Israel Institute of Technology, Haifa 3200003, Israel for their help in the Cryo-EM imaging and analysis.

## Author Contributions

Conceptualization: EK, YL, UE, EM, DP, OC.

Investigation: EK, YL, UE, HC, IHH, MA, AE, EBH, YD, SR, GSN, NA, MG, OC.

Funding acquisition: DP, EM and OC.

Supervision: OC, EM, DP.

Writing – original draft: EK, YL, UE, EM, DP, OC.

## Supplementary Materials

### Materials and Methods

#### Materials

Cholesterol, distearoyl-sn-glycero-3-phosphocholine (DSPC), and dimyristoyl-rac-glycero-3-methoxypolyethylene glycol (PEG-DMG) were from Avanti lipids (Alabaster, AL, USA). The proprietary ionizable lipid (Lipid 14) was synthesized in-house by Dr. Gonna Somu Naidu, as previously described (*27*). Custom and non-custom mRNA sequences and N1-Methyl-Pseudouridine modified nucleotides were from Trilink (San Diego, CA, USA) or synthesized in house by an IVT reaction using the MEGAscript™ T7 transcription kit and cleaned by the MEGAclear™ transcription clean-up kit both from Thermo Fisher Scientific (Waltham, MA, USA).

#### Design of mRNA constructs

Three mRNA constructs encoding the circular permutated F1 (Caf1) antigen (devoid of the *caf1* secretion signal and starting from P16 as described previously (*16*)). were designed as follows: SP-*caf1* encodes for the F1 protein, with the addition of a 5’ signal peptide (SP) sequence originating from the Ig light chain variable region (GenBank Acc. Number AAB01801(. ΔSP-*caf1* encodes for the F1 protein devoid of the SP sequence. SP-*caf1*-hFc is comprised of the SP-*caf1* sequence, followed by the sequence coding for the constant region of the human IgG1 (adopted from Genbank Acc. Number AEV43323.1). Luc control mRNA encodes the firefly luciferase (Genbank Acc. Number AB762768). All mRNA constructs included an initiator methionine, a Kozak consensus sequence, and were codon optimized for expression in mice.

#### LNP Preparation and Characterization

Ionizable lipid, Cholesterol, DSPC, and PEG-DMG were mixed at a molar ratio of 40:47.5:10.5:2 with absolute ethanol in a tube. mRNA payloads were suspended in 50mM Citrate buffer, pH 4.5. To create LNPs, a dual syringe pump was used to transport the two solutions through the NanoAssembler™ micromixer from Precision NanoSystem (Vancouver, British Columbia, Canada) at a total flow rate of 12 mL/min. The particles were then transferred into dialysis overnight against PBS. Particles in PBS were analyzed for size and uniformity by dynamic light scattering (DLS). Zeta potential was determined using the Malvern™ zeta-sizer (Malvern, Worcesrershire, UK). mRNA encapsulation in LNPs was calculated according to Quant-iT™ RiboGreen™ RNA Assay Kit (Thermo Fisher, Waltham, MA, USA), by calculating the percentage encapsulation at 100% - (mRNA-LNPs/mRNA-LNPs with triton). All mRNA-LNP formulations exhibited an encapsulation efficiency of >92%.

#### Cell lines

HeLa cells (ATCC CCL-2) were maintained at 37°C, 5% CO2 in Dulbecco’s modified Eagle’s medium (DMEM) supplemented with 10% fetal bovine serum (FBS), MEM non-essential amino acids (NEAA), 2 mML-glutamine, 1 mM sodium pyruvate, 100 Units/ml penicillin and 0.1 mg/ml streptomycin (P/S) (Biological Industries, Israel).

#### Cell transfection and Western blotting

One day before the transfection, HeLa cells were seeded in 24-well plates at a density of 10^5^ cells/well. At the day of the transfection, SP-*caf1*, ΔSP-*caf1* or SP-*caf1*-hFc mRNA-LNPs were added to the wells, and cells were harvested at 24-96 hours for detection of F1 protein in cell lysate (cellular fraction) and cell supernatant (secreted fraction). For cellular fraction evaluation, cells were washed with phosphate-buffered saline (PBS) and lysed with RIPA lysis buffer (Merck). The concentration of protein was measured by Bradford method, and equal amounts of protein were resuspended in protein sample buffer with beta-mercapthoethanol and boiled prior to separation. 30µg of cell lysate in 30µl buffer or 30µl of cell supernatant in buffer were separated by 12% SDS-PAGE. Gels were blotted by dry transfer with the iBlot™ Gel transfer device (ThermoFisher) to iBlot™ mini nitrocellulose membranes. Membrane was blocked for 1 hour at room temperature with 5% skim milk (DIFCO) in PBST. F1 expression was detected by overnight incubation with purified IgG fraction from serum of rabbit immunized with the recombinant F1 protein (1:10,000 dilution), followed by incubation with a secondary HRP-conjugated anti-rabbit antibody (Jackson Immunoresearch, 1:5,000 dilution) for 1 hour at Room temperature. Reactive bands were detected by development with SuperSignal™ West Pico PLUS Chemiluminescent Substrate (ThermoFisher) and detection by Amersham ImageQuant 800™ western blot imaging system (Cytiva life sciences).

#### Anti-F1 ELISA

Individual IgG titers generated against F1 were determined by ELISA as previously reported (*23*). Briefly, microtiter plates were coated with 500 ng of purified rF1 (provided by the Biotechnology Department at IIBR). Tested sera were serially diluted in 2-fold dilutions in a final volume of 50 μl and were incubated in the wells for 1 h at 37°C. Alkaline phosphatase-conjugated goat anti-rabbit IgG (1/2,000 dilution, Sigma) was used as the 2nd layer for rabbit anti-F1 IgG titer determination. Titers were defined as the reciprocal values of the endpoint serum dilutions that displayed OD405 values 2-fold higher than the normal serum controls.

#### Cryogenic Electron Microscopy (cryo-EM)

Samples were prepared in a closed chamber at a controlled temperature and at water saturation. A 5-6 µl drop of each suspension was placed on a 200-mesh transmission electron microscope (TEM) copper grid covered with a perforated carbon film. The drop was blotted, and the sample was plunged into liquid ethane (−183°C) to form a vitrified specimen, then transferred to liquid nitrogen (–196°C) for storage. Vitrified specimens were examined at temperatures below −175°C in a Talos F200C with field emission gun operated at 200Kv or a Tecnai T12 G2 TEM (FEI, Netherlands). Images were recorded on a Cooled Falcon IIIEC (FEI) Direct Detection Device by TIA software attached to the Talos or a Gatan MultiScan 791 camera by Digital Micrograph software (Gatan, U.K.) on the Tecnai. Volta PhasePlate (VPP) was used for contrast enhancement. Images are recorded in the low-dose imaging mode to minimize beam exposure and electron-beam radiation damage using lab procedures.

#### Ethics statement

This study was carried out in strict accordance with the recommendations for the Care and Use of Laboratory Animals of the National Institute of Health. All animal experiments were performed in accordance with Israeli law and were approved by the Ethics Committee for animal experiments at the Israel Institute for Biological Research (Permit Numbers M-55-21 and M-14-22). During the experiments, the mice were monitored daily. Humane endpoints were used in our survival studies. Mice exhibiting loss of the righting reflex were euthanized by cervical dislocation.

#### Animals

Female C57BL/6 mice (6–8 weeks old) were obtained from Envigo and randomly assigned into cages in groups of 10 animals. The mice were allowed free access to water and rodent diet (Harlan, Israel).

#### Animal vaccination experiments

Female C57BL/6 mice (6-8) weeks old were vaccinated intramuscularly (50ul to each hind leg muscle, for a total of 100ul) thrice with 5ug of SP-*caf1*, ΔSP-*caf1*, SP-*caf1*-hFc mRNA-LNPs or luciferase mRNA-LNPs (negative control). Alum-adsorbed recombinant F1 protein (80ug/mouse/dose) and alum (0.36% final conc./mouse/dose) were intramuscularly administered as positive and negative controls, respectively. The first vaccination study included the following groups: SP-*caf1* mRNA-LNPs (n=3), rF1+alum (n=3) and luciferase mRNA-LNPs (n=3). The second vaccination study included the following groups: ΔSP-*caf1* mRNA-LNPs (n=12), SP-*caf1*-hFc mRNA-LNPs (n=12), rF1+alum (n=6), luciferase mRNA-LNPs (n=8) and alum only (n=4). Blood samples were collected from the tail vein one day before boost administration. Spleens and blood samples were collected two weeks after administration of the last boost shot for evaluation of immunological responses.

#### Murine IFNγ ELISpot Assay

Mice spleens were dissociated in GentleMACS C-tubes (Miltenyl Biotec), filtered, treated with Red Blood Cell Lysing Buffer (Sigma-Aldrich, No. R7757), and washed. Pellets were resuspended in 1 mL of CTL-Test Medium (CTL, No. CTLT 005) supplemented with 1% fresh glutamine and 1 mM Pen/Strep (Biological Industries, Israel), and single cell suspensions were seeded into 96-well, high-protein-binding, PVDF filter plates at 300,000-400,000 cells/well. Mice were tested individually in duplicates by stimulation with a 15-mer peptide library spanning the F1 protein (10 μg/mL) (Genscript), Concanavalin A (Sigma-Aldrich, No. 0412; 2 μg/mL) as positive control, or CTL medium as negative control (no antigen). Cells were incubated with antigens for 24 h, and the frequency of IFNγ-secreting cells was determined using Murine IFNγ Single-Color Enzymatic ELISPOT kit (CTL, No. MIFNG 1M/5) with strict adherence to the manufacturer’s instructions. Spot forming units (SFUs) were counted using an automated ELISpot counter (Cellular Technology Ltd.).

#### Mouse infection

The fully virulent *Y. pestis* strain Kimberly53 (Kim53) was grown on brain-heart infused (BHI) agar plates at 28ºC. Several isolated and typical colonies were suspended in saline to generate a bacterial suspension at an optical density (660 nm) of 0.1 (equals to 108 cfu/ml). This bacterial suspension was serially diluted, and a dose of 100 cfu (100 LD_50_) was injected subcutaneously (s.c.) into the lower right backs of the mice. The infectious dose was verified by plating diluted bacterial suspensions onto BHI agar plates. Immunizations groups were as follows: ΔSP-*caf1* mRNA-LNPs (n=8), SP-*caf1*-hFc mRNA-LNPs (n=8), rF1+alum (n=6), luciferase mRNA-LNPs (n=4), alum only (n=4). Survival of infected mice was daily monitored for 21 days.

#### Statistical analysis

All values are presented as mean plus standard error of the mean (SEM). Statistical analysis was performed using either a two-way ANOVA with Tukey’s multiple comparisons test (for ELISA data) or a one-way ANOVA followed by post hoc Newman–Keuls test (for ELISA and ELISpot data) or log-rank (Mantel–Cox) test (for survival data) (*, p < 0.05; **, p < 0.01;, *** p < 0.001; ****, p<0.0001). All statistical analyses were performed using GraphPad Prism 8 statistical software.

